# The role of plasmids in the gut microbiome during the first year of life

**DOI:** 10.1101/2023.04.05.535656

**Authors:** Wanli He, Jakob Russel, Franziska Klincke, Joseph Nesme, Søren Johannes Sørensen

## Abstract

Plasmids are extrachromosomal self-replicating genetic elements that play a key role in bacterial ecology and evolution by shuttling diverse host-beneficial traits between bacteria. However, our understanding of plasmids is still limited, particularly in the human gut microbiota, and little is known about how they are acquired and become established in infants. In this study, we explored a longitudinal fecal metagenomic dataset obtained from 98 Swedish children who were followed during their first year of life. For this, we developed a bioinformatics pipeline for the complete sequence assembly and annotation of plasmids, together with the identification of plasmid contigs. We found that gut plasmids in these children were extremely diverse, particularly in the first four months of life, and this diversity decreased with maturation of the gut microbiota. Members of genera *Bacteroides* and *Bifidobacterium* were identified as major hosts of transmissible plasmids and important hubs of horizontal gene transfer in the early human gut microbiota. Additionally, we discovered that plasmids played a substantial role in expanding the gene repertoires of their bacterial hosts: approximately a quarter of unannotated plasmid genes were found only on plasmids and not on chromosomes. Together, our results provide the first characterization of the early acquisition and development of plasmids in the infant gut microbiome. Their diversity and abundance in the first months of life could benefit a variable and rapidly proliferating microbiota by providing increased adaptability in a highly competitive environment.

## Introduction

Plasmids are self-replicating extrachromosomal genetic elements that are broadly present in bacteria and archaea. These molecules show remarkable diversity in size^1^, copy number^2^, GC content^3^, replication mechanism^4^, transmission mode^5,6^, DNA topology (circular or linear)^7^, genetic cargo, and host range^8^, among other features. Importantly, plasmids carry both backbone or ‘core’ genes that are instrumental to their vertical and horizontal transmission and self-replication^5,9,10,11^ as well as genetic cargo that codes for genes involved in virulence, ecological interactions, anti-phage systems, and antibiotic resistance, and many unknown functions^12,13,14,7^.

Plasmids play a key role in bacterial ecology and evolution, especially via horizontal gene transfer (HGT). Plasmids are key drivers of HGT and can be transferred at high rates through a variety of mechanisms, including conjugation (including plasmid mobilization and conduction), transduction, transformation, and vesiduction^5,15,16,17^. As a result, beneficial traits are rapidly transferred within and between species of bacteria, eventually contributing to increased host fitness^18^. In addition, high copy numbers of plasmids cause gene dosage effects that increase gene expression^19^ as well as gene variability through gene maturation and recombination^20,21^. However, plasmids also impose a burden on their bacterial host from the demands of the plasmid life cycle (for example, plasmid conjugation^22^, replication, and gene expression^23^) and from conflicts between chromosomal-encoded and plasmid-encoded proteins within the host bacterium^24^.

Studies on plasmids have historically focused on single bacterial isolates or mathematical modeling. For example, researchers have investigated plasmid traits in specific bacteria^25,13^, studied plasmid-mediated antibiotic resistance genes (ARG)^26,27^, and performed theoretical studies of plasmid persistence and dynamics^12,28^. Recently, developments in high-throughput metagenomic sequencing have made it possible to study the plasmid metagenome, known as the plasmidome, and some studies have used this approach to explore plasmid community in the gut environment^29,30,31^. However, our understanding of plasmids is still limited by methodological challenges, including the reliable identification of plasmids and the detection of their individual bacterial hosts in a metagenomic catalog.

Moreover, we lack knowledge about the biology of the human plasmidome beyond archetypal pathogen-associated plasmids. In particular, there has been no comprehensive assessment of the dynamics of the gut plasmidome in early life, even though numerous studies in the last decade have emphasized the importance of the gut microbiome for healthy infant development^32,33,34^ and plasmids are known to improve the fitness and environmental adaptability of their bacterial hosts^35,13^. It is plausible that plasmid assemblages in the human gut could shape the colonization and development of bacterial communities during microbiome establishment and maturation. As a starting point, what is needed to test this hypothesis is an overview of the gut plasmidome and plasmid hosts in the human gut that characterizes how they interact, coexist, and develop.

To study the gut plasmid community in early life, we developed a workflow (https://github.com/Wanli-HE/Plaspline) that systematically integrates available tools for plasmid assembly and identification. We then used this pipeline to build an extensive plasmid catalog from a longitudinal gut metagenomic dataset obtained from 98 Swedish children followed during their first year of life (Figure 1). We were able to recover thousands of novel circular plasmid sequences with no close relatives in current reference databases, considerably expanding our knowledge of the genetic diversity of plasmids. By analyzing the replication backbone genes of these plasmids, we reconstructed the phylogenetic relationships in the gut plasmid community and identified 328 distinct replicon groups (Figure 2). We were also able to link plasmids to their most likely bacterial hosts using plasmid sequence similarity networks, revealing that most plasmids in early life gut are likely harbored by species belonging to *Escherichia* and *Klebsiella* genus, which is expected based on previous knowledge, but also in large part by species of genus *Bacteroides* and *Bifidobacterium* that are not well represented in reference plasmid databases (Figure 3). This study provides the first overview of the gut plasmidome in early life, revealing the abundance and diversity of plasmids in the human gut and shedding light on their potential roles in the gut microbiota of infants.

**Figure 1.**
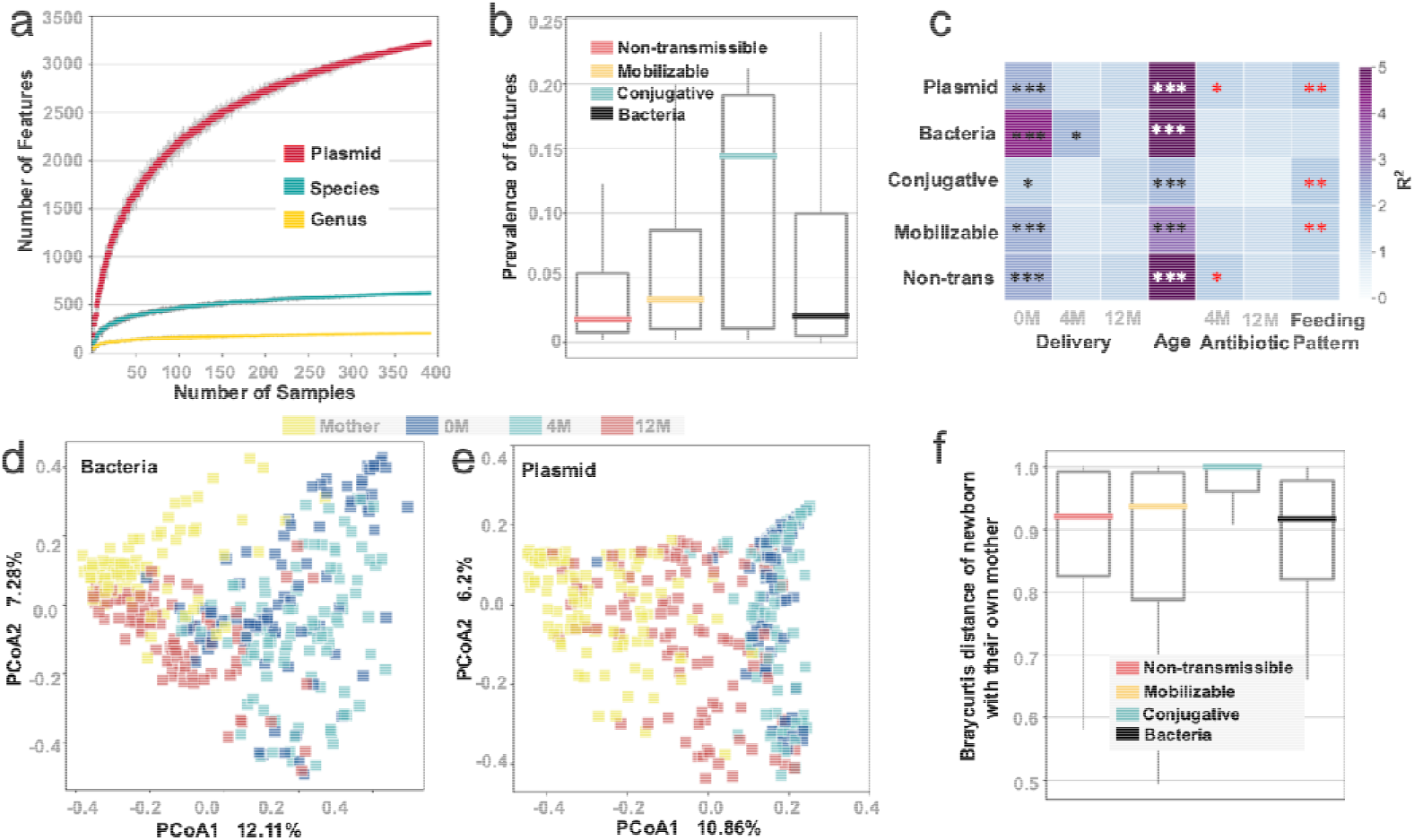
**a**: Number of bacterial taxa and plasmids discovered as a function of sample size. Color bars denote mean values. Error bars display SD of 30 resamplings. **b**: Prevalence of different types of plasmids and bacteria among all samples (392). Bacterial prevalence calculated at the species level; plasmids are grouped according to their mode of mobility. **c**: Factors that affect the composition of bacterial and plasmid assemblages, as determined by PERMANOVA analysis of Bray-Curtis distances; stars indicate statistical significance (*P ≤ 0.05, **P ≤ 0.01, ***P ≤ 0.001) and color indicates the value of the test statistic R2. **d and e**: Visualizations of principal coordinates analyses (PCoA) based on Bray-Curtis distances representing the beta-diversity of the bacterial (d) and plasmid (e) communities. **f**: Bray-Curtis distances between the assemblages (different types of plasmids and bacteria) in infants and their respective mothers.

**Figure 2.**
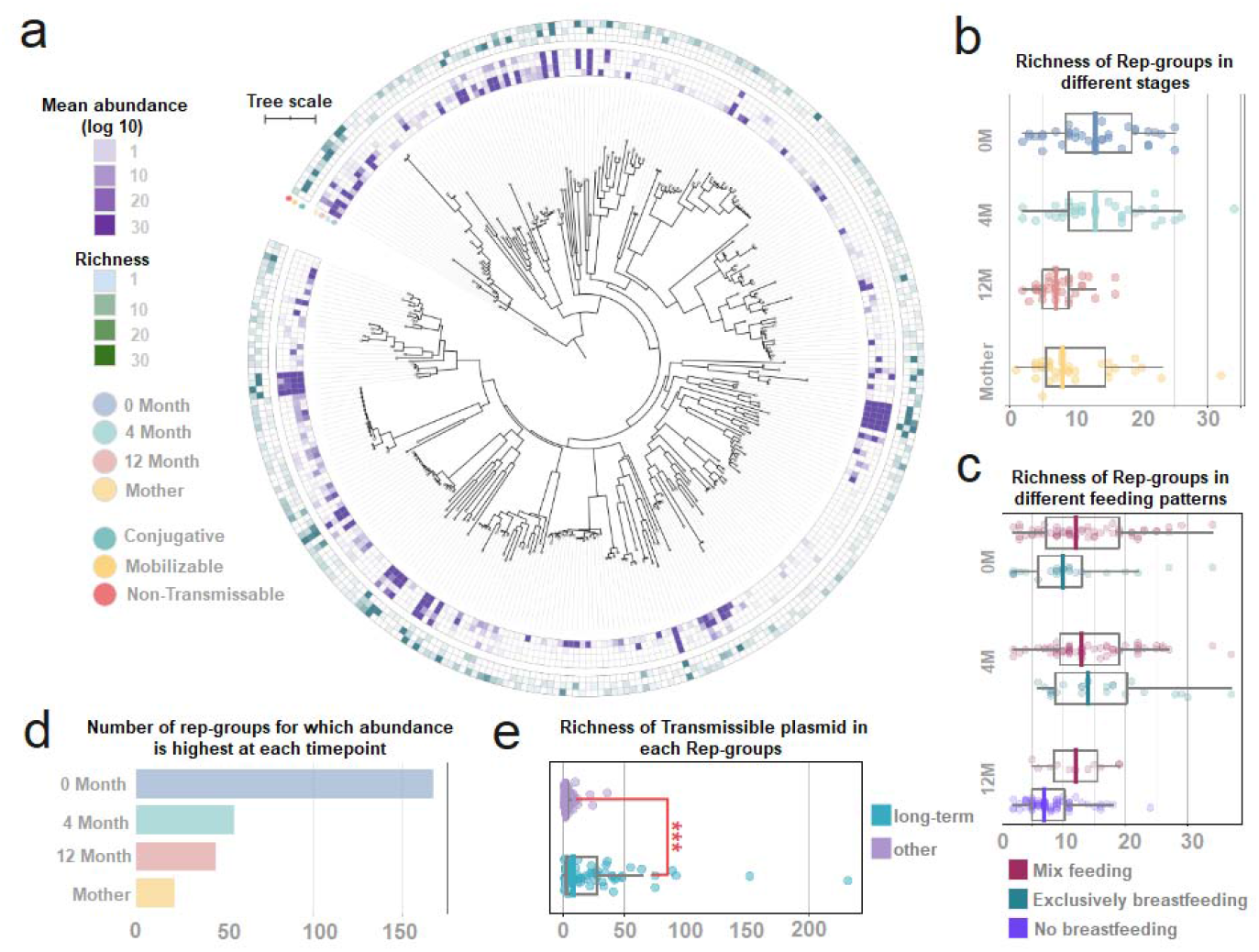
**a**: A phylogenetic tree of gut plasmids was constructed based on their replicase genes, from which 328 rep-groups were identified. The inner layer of the heatmap surrounding the tree depicts the abundance of each plasmid in samples of different ages, from newborn to mother (inner to outer); the outer layer represents the richness of plasmids from each mobility group (conjugative, mobilizable, or non-transmissible) in each rep-group. **b**: The richness of rep-groups in samples from all ages of vaginally born children and their mothers. **c**: The richness of rep-groups in samples from children with different breastfeeding patterns. **d**: The number of rep-groups whose abundance was highest at a given timepoint. **e**: The richness of transmissible plasmids in rep-groups that were found in children at all timepoints (long-term) versus rep-groups that were not.

**Figure 3.**
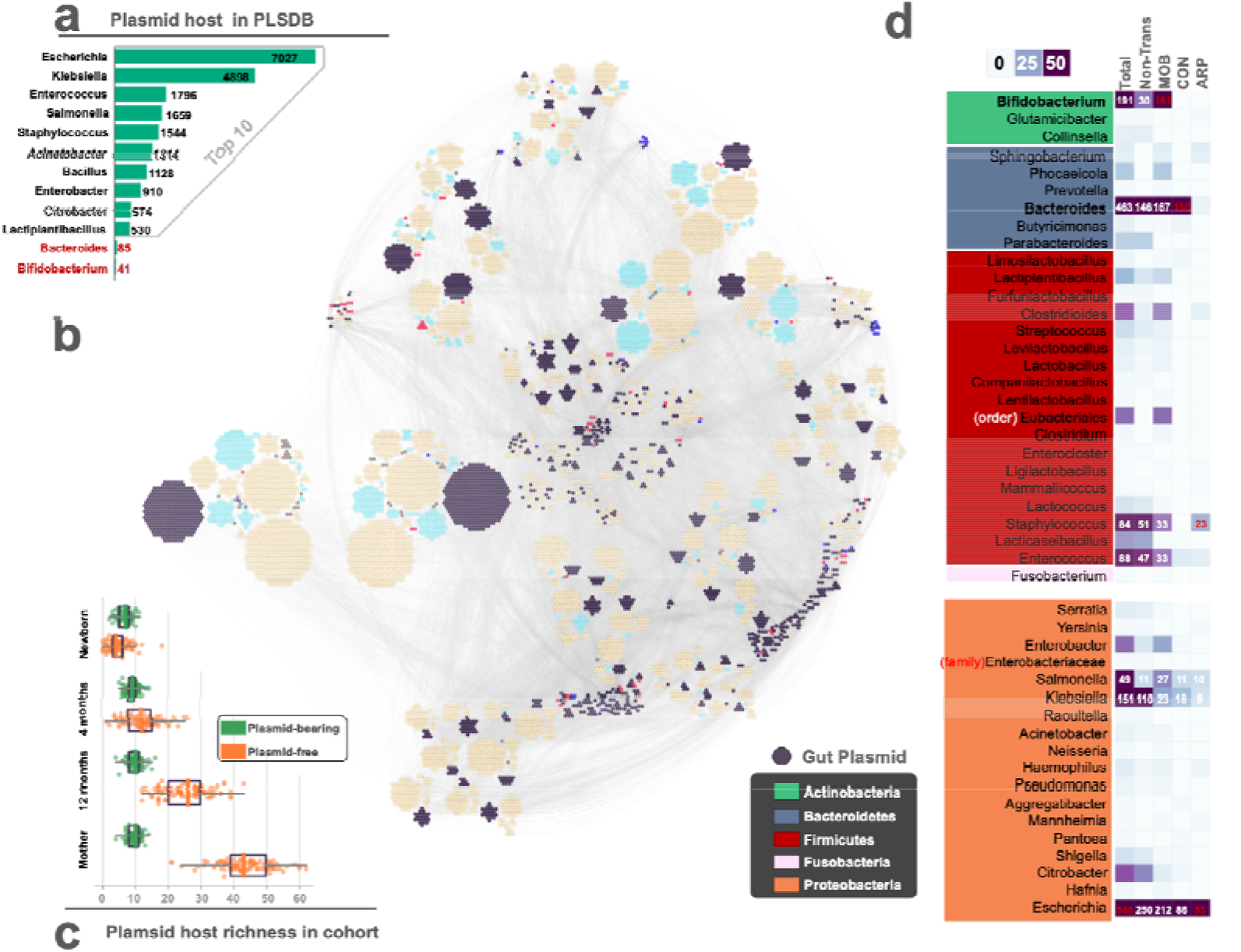
**a**: Numbers of plasmids associated with the top 10 plasmid host genera in PLSDB (2021-06-23) plus *Bacteroides* and *Bifidobacterium*. **b**: Associations between gut plasmids in this study and plasmids in PLSDB based on JI distances; only edges for which the JI distance >0.3 are displayed. Nodes representing PLSDB plasmids are colored by host phylum; nodes representing our plasmid catalog are colored in black. **c**. Richness of plasmid-linked bacterial host species compared to unlinked bacterial species among children of different ages and their mothers. **d**. Richness of plasmids found in different bacterial host genera. ‘Total’ indicates richness of all plasmids; ‘Non-trans’, ‘MOB’, ‘CON’, and ‘ARP’ indicate non-transmissible, mobilizable, conjugative, and antibiotic resistance plasmids, respectively.

## Methods

### Study population and sampling

The metagenomic data were obtained from a Swedish cohort of 98 children and their mothers; the children were sampled at three timepoints and the mothers were sampled once. The study population, sampling scheme, collection of metadata, and sequencing protocols are described in Bäckhed et al.^36^. Sequencing reads were downloaded from SRA (ERP005989). All infants were included in our analysis (described below) of the effect of mode of delivery on plasmid abundance and diversity (n=83 delivered vaginally, n=15 delivered by C-section).

### Reconstruction of plasmid catalog

The gut plasmid catalog was constructed using *Plaspline* (version: v1.2) (https://github.com/Wanli-HE/Plaspline.git). First, *de novo* metagenome assembly was performed using meta*SPAdes*^37^, with sequence reads (trimmed and filtered) as the input and default parameters applied (*k-mer* 21,33,55) for each sample. Contigs longer than 1 kb were selected for downstream analysis. Next, circular contigs were reconstructed by *metaSPAdes* (version 3.14.1) using the “*-plasmid*” option^38^ and *SCAPP* (version: v0.1.4)^39^ with max *k-mer* 55. To remove non-plasmid circular elements from the set of circular contigs, we used *plasmidVerify* (https://github.com/ablab/plasmidVerify - version: release May 1, 2020), which classifies contigs as plasmid or non-plasmid based on their gene content. A non-redundant catalog of circular plasmid genomes was constructed using *mmseqs* (version: 12.113e3)^40^ with sequence identity >0.9 and coverage >0.95 (“--cov-mode” 0). To obtain a more comprehensive catalog of plasmids, particularly at the gene level, the next step of the *Plaspline* workflow utilized *Plasforest* (version: v1.3)^41^ and *Platon* (version: 1.4.0)^42^ to detect plasmid fragments from all assembled contigs. After merging the results of these two tools, a non-redundant draft catalog (linear plasmid contigs) was generated using *mmseqs* (version: 12.113e3) with sequence identity >0.85 and coverage >0.8 (“--cov-mode” 2). Finally, the draft catalog was compared to the circular catalog using *BLAST* (version: 2.9.0+), and any contigs in the draft catalog that matched those in the circular catalog were removed (identity >0.85, coverage >0.8). The circular and the draft catalogs were then merged to create the final plasmid catalog.

### Plasmid genome and gene abundance calculations

To calculate the abundance of plasmid genomes and plasmid genes, *Plaspline* uses *Bwa* (version: 0.7.17)^43^, *Samtools* (version: 1.9)^44^, and *Msamtools* (https://github.com/arumugamlab/msamtools - version: 0.9). First, reads were filtered based on mapped length (>60bp), identity (90%), and read coverage (80%). Then, contigs/genomes with less than 55% mapped coverage were removed. Finally, abundance was calculated and normalized based on the number of fragments per kilobase of sequence mapped per million reads (FPKM). A gene’s abundance was calculated as the sum of the abundance of all contigs/genomes that contained that gene.

### Annotation of plasmid catalog

*Plaspline* makes use of the *MOB-recon* and *MOB-typer* modules from *MOB-suite* (version: 2.0.1)^45^ software to classify plasmids. These modules classify putative plasmids from metagenome contigs by searching for replication genes (rep), mobilization proteins (relaxase), genes encoding the mate-pair formation system (MPF), and origin of transfer (oriT) sites. Gut plasmids are then annotated based on the presence or absence of these plasmid markers.

### Plasmid phylogenetic tree

A plasmid phylogenetic tree was constructed using plasmid replicon marker genes. MAFFT (version: v7.475)^46^ was used to perform multiple alignment of replicase protein sequences. The multiple alignment results were then fed into IQ-TREE (version: 2.0.3)^47^ to reconstruct a phylogeny of plasmids, with the parameters *“-MFP”* to determine the best-fit model of the SH-like approximate likelihood ratio test using 1000 bootstrap replicates. The final phylogenetic tree was visualized and annotated using iTOL (v4)^48^.

### Plasmid host linking

The most-probable host of each plasmid reconstructed from the gut metagenomes was inferred based on genomic similarity between the plasmid sequences assembled in this study and known plasmids from the PLSDB database (version: 2021-06-23), which in most cases includes information on the host from which the plasmid was isolated^49^. We estimated genomic similarity by calculating the Jaccard Index (JI) based on shared 21-bp k-mers, with a minimum JI threshold of 0.3, as described in Acman et al.^50^. JI values were calculated using *Bindash* (version 1.0)^51^. If a plasmid matched with more than one host in PLSDB, all matching hosts were recorded. Finally, the bacterial host was annotated according to its linked plasmid host information in PLSDB.

### Taxonomic annotation and calculation of abundance

Taxonomic profiling of reads was carried out using *MetaPhlAn* (version: 3.0)^52^ at the species level.

### Gene annotation

Gene functions were annotated using eggNOG-mapper (version: 2.1.7)^53^. Virulence factors were annotated based on information in the VFDB database (version: before JUN 2022)^54^. Antibiotic resistance genes were annotated using the CARD Resistance Gene Identifier (RGI, version: v5.2.1)^55^.

### HGT network construction

Based on the number and identity of plasmids associated with different hosts, we constructed a network of putative horizontal gene transmission events. Specifically, an HGT event was presumed to have occurred if plasmids with the same replication gene were found in two or more bacterial hosts. We then refined the HGT networks to examine children at different ages and with different feeding patterns and mode of delivery (Figure 4a; Supplementary figure 1a). In these analyses, the number of HGT events, and the number of non-transmissible and transmissible plasmids that were involved in HGT events, was calculated in each individual and then compared between different groups (Figure 4b, Supplementary figure 1b).

**Figure 4.**
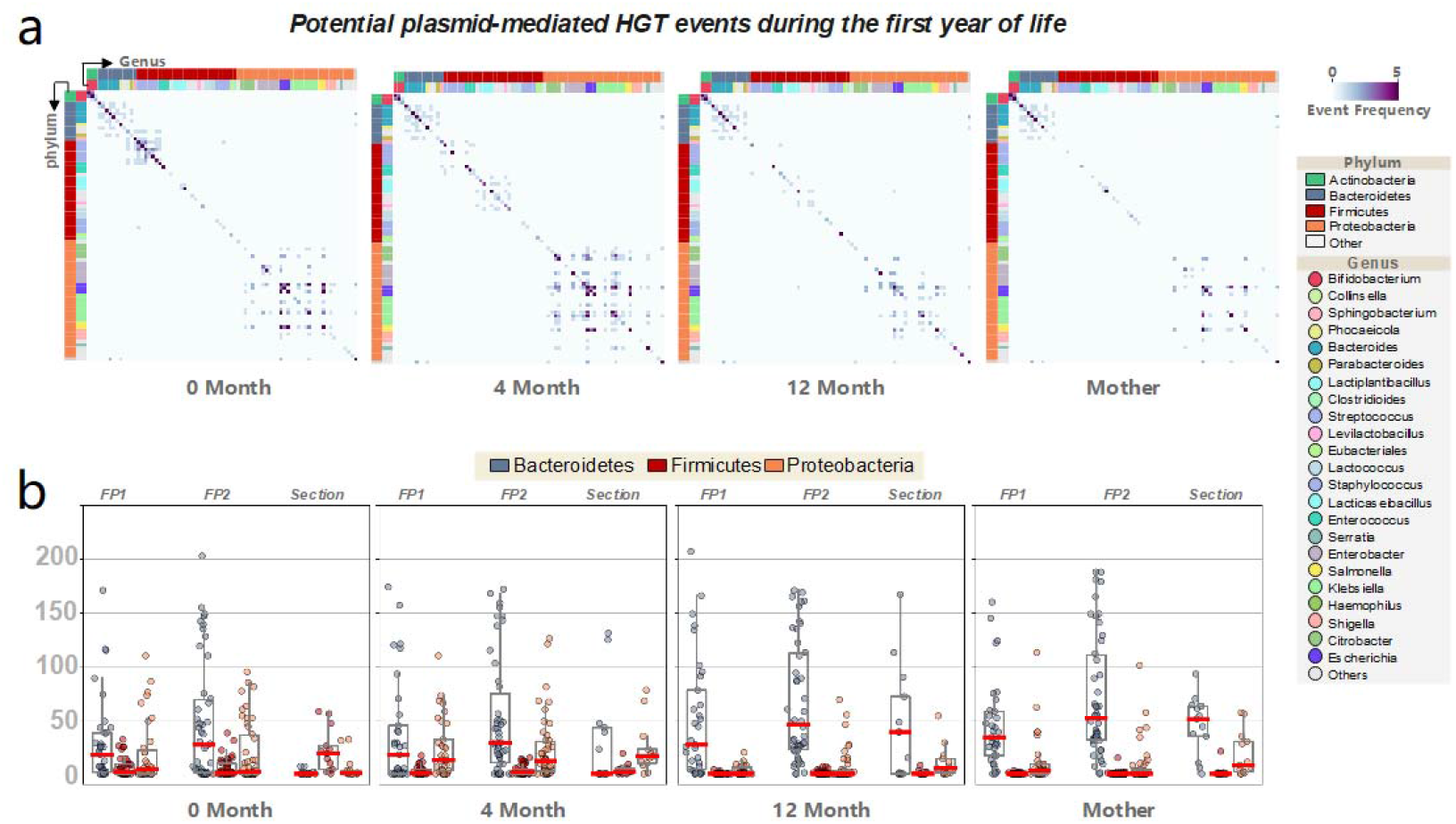
**a**: Networks of potential plasmid-mediated horizontal gene transfer (HGT) between species were constructed for all children and mothers. The inner strip represents genus-level annotations, and the outer strip depicts phylum-level assignments. **b**: The HGT events in each child and mother were compared based on patterns of breastfeeding and mode of delivery. ‘FP1’ represents children who were exclusively breastfed in the first four months but received no breastfeeding at 12 months; ‘FP2’ represents children who received breastmilk and formula (mixed feeding) in the first four months and any amount of breastfeeding at 12 months of age. Children born by C-section (labeled ‘section’) were considered separately given the well-known impact of delivery mode on establishment of the gut microbiome.

### Statistical analysis

All statistical analyses were performed using python (3.7.6) with the packages “*scipy*.*stats* (1.7.3)” and “*skbio* (0.5.6)”. Alpha and beta diversity of plasmid and bacterial communities were calculated using the functions “*skbio*.*diversity*.*alpha_diversity*” and “*skbio*.*diversity*.*beta_diversity*”.

To estimate the prevalence of different types of plasmids relative to that of bacteria, Wilcoxon rank-sum tests were performed using “*skbio*.*stats*.*ranksums*”. We explored the effects of four factors (age, feeding pattern, delivery mode, and antibiotic usage) on the composition of the gut plasmidome by examining correlations between the composition of plasmid assemblages and that of microbial communities (Bray-Curtis distance); this was carried out using permutational multivariate analysis of variance (PERMANOVA) with the functions “*skbio*.*stats*.*ordination*.*pcoa*” and “*skbio*.*stats*.*distance*.*permanova*” (permutations = 999). To assess how community composition changed with age, we paired the plasmid and bacterial assemblages of each child and performed Wilcoxon signed-rank tests using “*skbio*.*stats*.*wilcoxon*”.

To compare plasmid richness (as assessed by differences in their replication backbone genes) between children at different ages and with different feeding patterns, we used Wilcoxon signed-rank tests and Wilcoxon rank-sum tests, respectively. We also used Wilcoxon rank-sum tests to compare the richness of transmissible plasmids in groups of persistent (present from birth to 12 months) versus transient plasmids. The phylogenetic influence on transmissibility was assessed with Pagels λ using the R package ‘*phytools*’, with the ratio of transmissible to non-transmissible plasmids in each replication-gene group as the continuous variable. If the number of non-transmissible plasmids was 0 it was reset to 1 to avoid infinite ratios.

We used Wilcoxon signed-rank tests to compare frequency and density of plasmid transfer events for the three main phyla with the highest number of potential plasmid HGT events as defined above. in the same group of children at different ages. The same method was used to analyze the richness of non-transmissible and transmissible plasmids.

To compare patterns of gene function and pathway coverage between plasmids and chromosomes, we paired them and used Wilcoxon signed-rank tests. Pathway coverage was calculated as the number of genes discovered from a given pathway divided by the total number of genes in the pathway, ARG richness was calculated at ARO level, virulence factors were assessed at the level of annotation identity, and defense systems were examined at the D-level from KEGG annotation. We then compared patterns of gene abundance (functional groups) between plasmids and chromosomes using Wilcoxon rank-sum tests after correction for the false discovery rate (python function ‘*fdrcorrection*’ from ‘*statsmodels*”).

## Results

### Plasmids are highly diverse in the early life humanx gut

Through longitudinal sampling of a cohort of 98 Swedish infants and their mothers^36^, 392 fecal samples were sequenced and processed using our *Plaspline* pipeline. From this, we were able to construct a catalog of the plasmids in these early life gut metagenomes that included, in total, 3,377 complete circular plasmids and 11,977 plasmid contigs (draft). Our sampling effort appeared to capture the majority of the bacterial diversity present in this population, with 83% of bacterial species detected with only 40% of the 392 samples (Figure 1a). In contrast, the number of plasmids discovered continued to increase dramatically with further sampling. This indicates that many rare plasmids remained undiscovered and highlights the exceptional diversity of these genetic elements. Non-transmissible plasmids—those with no known/detectable mobilization or conjugative genes—represented the largest group of identified gut plasmids (complete circular plasmid: 64.5%, total plasmid contigs: 88.4%), with conjugative (complete circular plasmid: 6.1%, total plasmid contigs: 2.2%) and mobilizable (complete circular plasmid: 29.4%, total plasmid contigs: 9.4%) plasmids constituting a clear minority of the population. However, the prevalence of non-transmissible plasmids was significantly lower than that of bacterial species (Wilcoxon rank-sum test; p=0.016), indicating a higher variability among samples in this type of plasmid than in bacterial species. In contrast, the prevalence of transmissible plasmids (mobilizable and conjugative) was significantly higher than that of bacterial species (Wilcoxon rank-sum test; mobilizable plasmid vs. bacteria, p=0.004; conjugative plasmid vs. bacteria, p<0.001) (Figure 1b), indicating that mobilizable and conjugative plasmids were more widely shared among individuals than bacterial species were.

Given the high degree of inter-individual differences among plasmid communities, we hypothesized that the plasmid assemblages might be more sensitive than the overall bacterial community (as described at the species level) to environmental factors. As expected, the age of the human host and mode of delivery (vaginal vs. Cesarean) strongly affected both bacterial and plasmid communities (Figure 1c). However, the species-level bacterial community was not significantly different between children with different breastfeeding patterns or histories of antibiotic use, whereas the plasmid communities were (Figure 1c). Interestingly, in 4-month-old children, a clear effect of mode of delivery was still detectable in the bacterial communities, but this effect had already faded in the plasmid community, which suggests that the latter assemblages may mature more quickly than the former.

When we performed pairwise comparisons between the children and their respective mothers, we found that the plasmid communities were more distinct than the bacterial communities were (Figure 1d and e) (Wilcoxon signed-rank test; p< 0.001). However, when we divided the plasmids into mobility groups, this difference was only seen for conjugative plasmids (Wilcoxon rank-sum test; bacteria vs. conjugative plasmid: p<0.001; bacteria vs. mobilizable plasmid: p=0.89; bacteria vs. non-mobilizable plasmid: p=0.16) (Figure 1f), suggesting that conjugative plasmids may colonize the infant gut earlier and faster than other plasmids. Moreover, the distance (dissimilarity) between the plasmid communities in newborns and 4-month-olds was significantly lower than that between the bacterial communities (Wilcoxon signed-rank test; p<0.001), and similar results were observed between 4-month-olds and 1-year-olds (Wilcoxon signed-rank test; p4m-12m=0.019). These results also support the hypothesis that the maturation of the plasmid community was faster than that of the bacterial community.

### Phylogenomic representation of the gut plasmidome in early life

To obtain a systematic measure of gut plasmid diversity, we classified each contig in the total plasmid repertoire (15,354 contigs) according to its replicon backbone genes, thus obtaining a total of 328 replicon groups (rep-groups). The plasmids within each rep-group were further characterized as conjugative, mobilizable, or non-transmissible based on identified backbone genes involved in mobility. We then calculated the abundance and richness of each replicon group in all samples obtained from our cohort, including the mothers. Of the 328 total rep-groups, 67.68% were most abundant in the first four months of life (Figures 2a and 2d), while only 6.71% were most abundant in the mothers (Figure 2d). The diversity of rep-groups was significantly higher in newborns and 4-month-olds compared to 12-month-olds and mothers (newborn vs. 12M, mother; Wilcoxon signed-rank test; p=1.67e-23, p=0.0016) (4M vs. 12M, mother; Wilcoxon signed-rank test; p=1.1e-09, p=3.2e-08). The highest diversity was found in the plasmid communities of 4-month-old children (newborn vs. 4M; Wilcoxon signed-rank test; p=0.003) (Figure 2b). Overall, these data clearly indicated that plasmids were enriched in the first four months of life.

Next, we examined the influence of feeding patterns on gut plasmid richness. We found no difference between mix-feeding and exclusive breastfeeding in newborn and 4-month-old children (Mix-feeding vs. exclusive breastfeeding; Wilcoxon rank-sum test; p_new_=0.12, p_4M_=0.58). However, plasmid richness was significantly lower in children with no breastfeeding at 12 months compared to those that were still mix-feeding (Wilcoxon rank-sum test; p=0.01) (Figure 2c), suggesting that breastfeeding may play an important role in shaping plasmid communities in early life.

The majority of plasmids were present in all sampling series, including the mothers (Figure 2a). Notably, we also found that the persistent rep-groups—those that were found in children at all sampled timepoints—contained a higher number of transmissible plasmids than more-transient rep-groups (Figure 2e; Wilcoxon rank-sum test; p=7.42e-10), suggesting that plasmid mobility may be important for their persistence. Furthermore, these higher-mobility rep-groups were found to be more clustered on the phylogenetic tree than would be expected by chance (phylogenetic signal, Pagel’s λ: 0.85; p-value (based on LR test): 2.01e-11).

### *Bacteroides* and *Bifidobacterium* are the main hosts of transmissible plasmids in the human gut

To investigate the relationship between plasmids and their bacterial hosts, we compared the draft and circular plasmids obtained here with published plasmids whose bacterial hosts are known (see methods). Using this approach, we were able to link 1,966 of 15,425 (12.7%) gut plasmids to a plasmid from PLSDB with a known host (2021-06-23). Many highlighted plasmids in PLSDB had *Escherichia* or *Klebsiella* as their predicted host, while only a small group of plasmids were from *Bacteroides* and *Bifidobacterium* (Figure 3b). This may primarily be due to database bias, since PLSDB contains thousands of plasmids from *Escherichia* and *Klebsiella*, but only 85 from *Bacteroides* and 41 from *Bifidobacterium* (Figure 3a). Numerous plasmids in our sample demonstrated similarity to published plasmids from *Bacteroides* and *Bifidobacterium*, indicating that these bacteria were major hosts for the infant gut plasmidome in this study. Furthermore, this suggests that many rare plasmids in these bacteria remain to be discovered and characterized.

Interestingly, many of the plasmids associated with *Bacteroides* and *Bifidobacterium* were transmissible; indeed, many conjugative plasmids were found to be associated with *Bacteroides*, even more so than with *Escherichia* (Figure 3d). However, *Escherichia* and *Staphylococcus* were linked with the highest number of antibiotic resistance plasmids (Figure 3d). This is in line with reports that these two bacteria are responsible for most fatalities linked to antimicrobial resistance worldwide, a feature that is often transferred via plasmids^56^.

We found that most plasmid replicon groups were shared by a limited number of bacterial taxa. Indeed, of the 328 replicon groups, only 5 were found in 4 different genera, while 9 rep-groups were present in 3 genera and 27 were present in 2 genera. In most of these cases, the host bacteria belonged to the same phylum (Supplementary Figure 1a and Supplementary Figure 2). Plasmids with a broad host range (i.e., two or more hosts, regardless of the phylogenetic distance between hosts) were most often found in Proteobacteria; specifically, plasmids in genus *Escherichia* were often shared by *Klebsiella* and *Salmonella*. Conversely, plasmids carried by Actinobacteria were only shared within a single host genus: *Bifidobacterium*. Different taxa of bacteria also differed in the specificity of their plasmids: for example, all rep-groups associated with order Eubacteriales and most rep-groups associated with genus *Citrobacter* were also found in other taxa. For plasmid rep-groups associated with *Escherichia*, 60% were shared with other genera.

Rep-groups found in *Bifidobacterium* were only found within that genus (Supplementary Figure 1a). Consistent with the analyses presented above, we also observed higher diversity and relative abundance of plasmids in the first 4 months (Supplementary Figure 1b), as shown by the plasmid dynamics in the genera *Bifidobacterium, Staphylococcus, Lacticaseibacillus, Citrobacter, Escherichia*, and *Klebsiella*. However, the abundance and diversity of plasmids carried by *Bacteroides* increased with children’s age, demonstrating how plasmid abundance and diversity were directly associated with bacterial host dynamics (Supplementary Figure 5).

**Supplementary figure 1.**
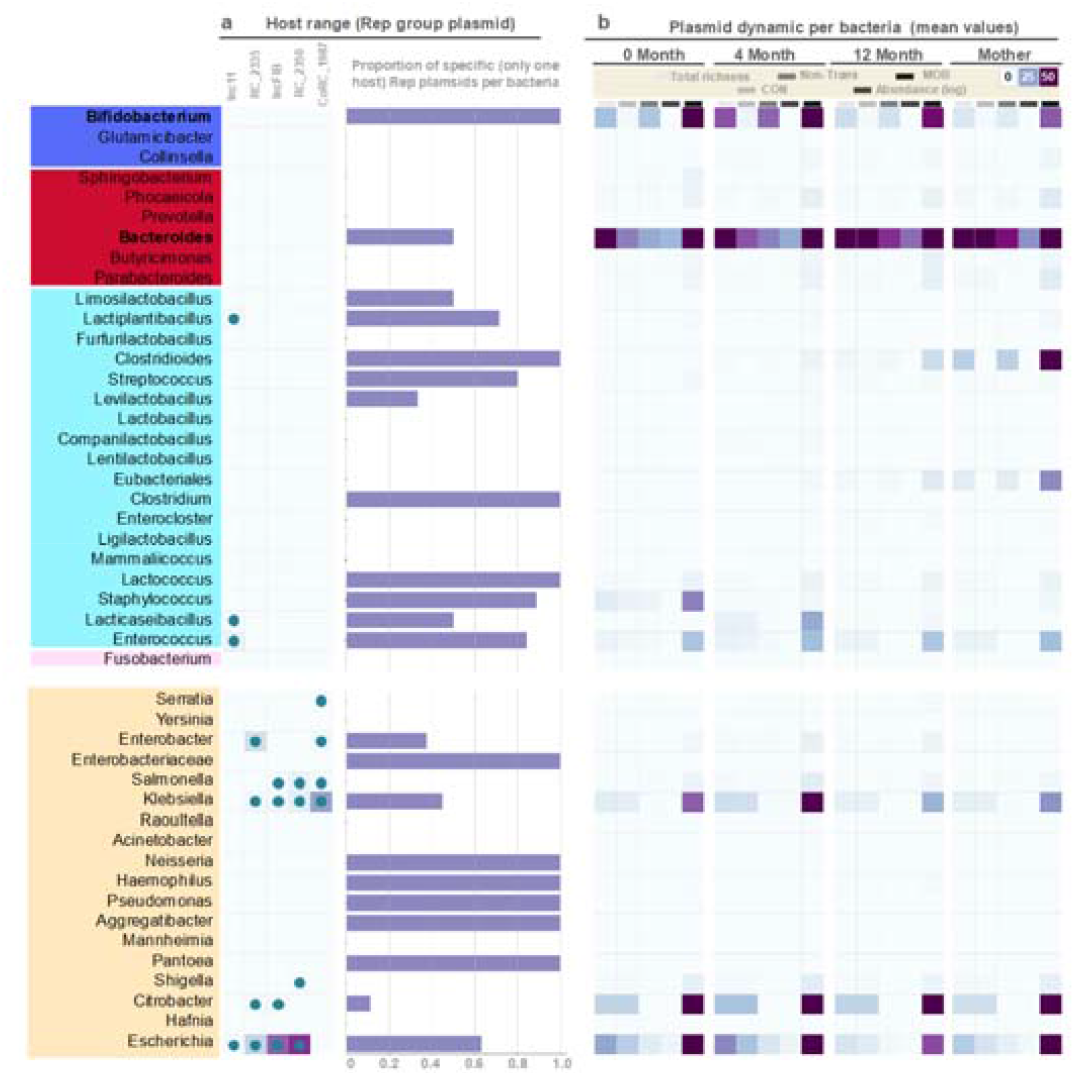
a: Heatmap depicting the richness of the rep-groups with the broadest bacterial host range in this study (four hosts for each rep-group). Bar plot shows the proportion of host-specific rep-groups found in each bacterium. b: Richness and abundance (FPKM) of bacterial plasmids (complete circular only) in children of different ages and their mothers.

**Supplementary figure 2.**
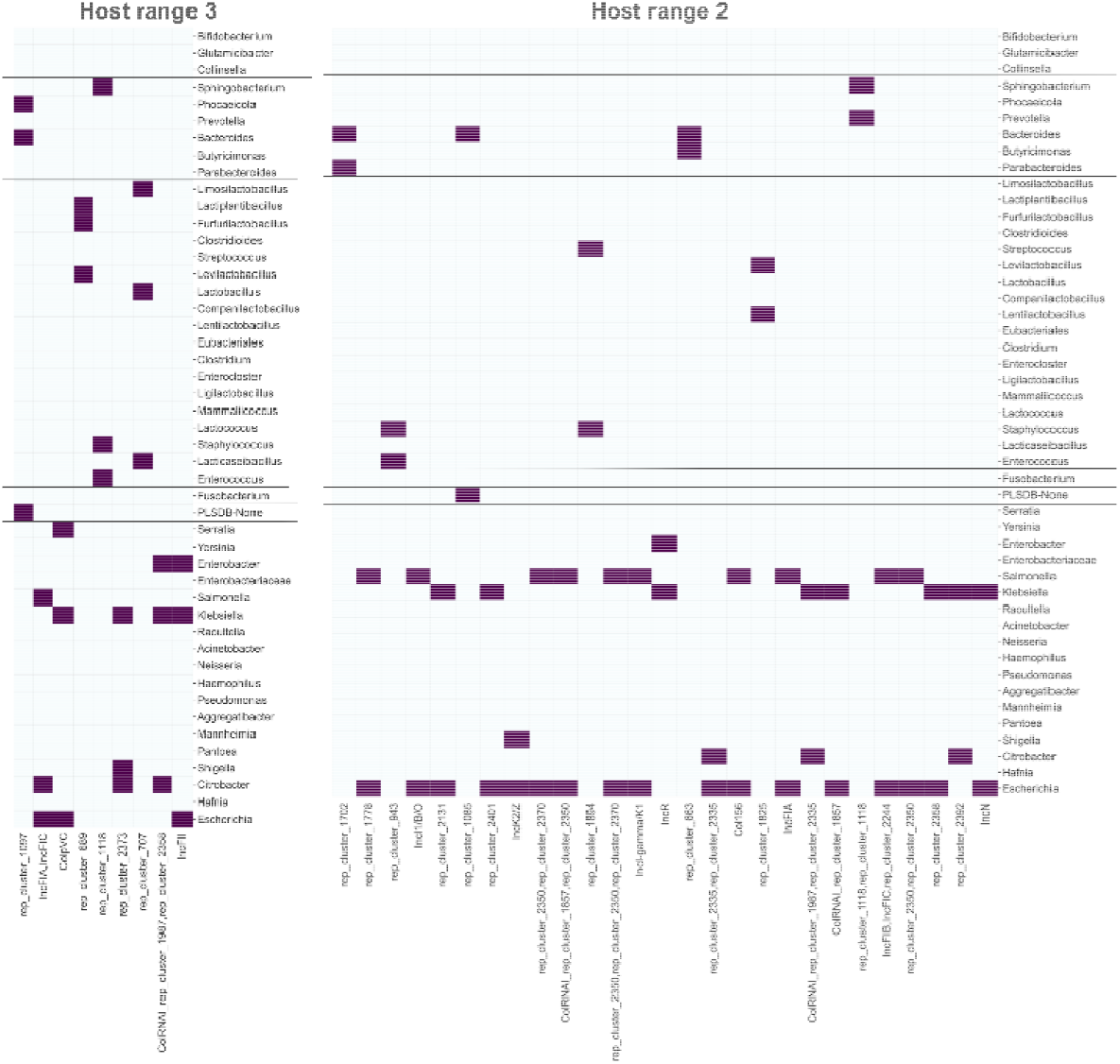
Heatmap depicting the richness of plasmid rep-groups groups found in three (left) or two (right) hosts

### The plasmid-mediated HGT network during the first year of life

**Supplementary figure 3.**
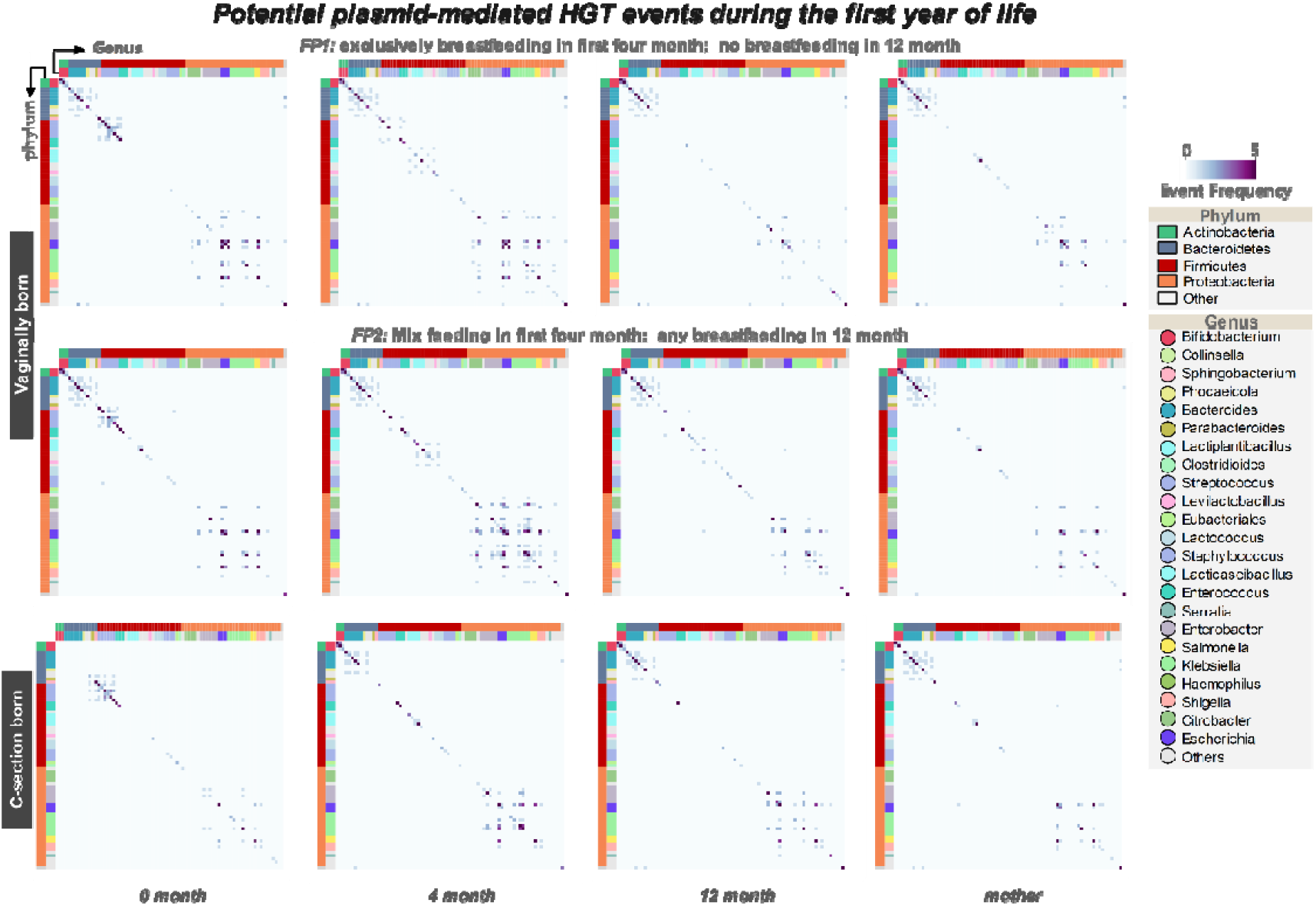
Networks of potential plasmid-mediated horizontal gene transfer (HGT) between bacteria in the gut microbiota of children at different ages. Data are presented based on the mode of delivery and breastfeeding patterns of children.

As plasmids are the main vectors of horizontal gene transfer (HGT) between bacteria, our next step was to analyze the potential plasmid-mediated connections between bacterial hosts in the early life gut microbiome. To construct the HGT network, we focused on the plasmids that were successfully linked to bacterial hosts in Figure 3b and grouped them according to their replication backbone genes (rep-group). Potential HGT events were defined as the detection of members of the same rep-group in different bacterial hosts. This information was used to construct HGT networks for each child and mother in the cohort; we then grouped children by their age, feeding pattern, and delivery mode to investigate the influence of these factors. We found that the majority of HGT events occurred in phyla Bacteroidetes, Firmicutes, and Proteobacteria, particularly in the first four months of life (Figure 4a, Supplementary Figure 3). Moreover, almost all HGT events took place within a single phylum (Figure 4a, Supplementary Figure 3), suggesting that most plasmid-mediated HGT events occurred between phylogenetically related bacteria.

Importantly, we observed the highest frequency of HGT in phylum Bacteroidetes; this was true for all ages of children and their mothers (Figure 4b; Wilcoxon signed-rank test, Supplementary Table 1 s1). This pattern became even more obvious after we normalized the network density, since Proteobacteria and Firmicutes had more species participating in HGT (Supplementary Figure 4; Wilcoxon signed-rank test, Supplementary Table 1 s2). To explore the question of why Bacteroidetes had such strong HGT networks, we examined the number of transmissible plasmids in each rep-group. We found that, indeed, members of phylum Bacteroidetes had the highest number of transmissible plasmids compared to other phyla, whereas Proteobacteria had more non-transmissible plasmids (Figure 4b, Supplementary Figure 4; Wilcoxon signed-rank test, Supplementary Table 1 s3 and s4). Taken together, these data indicate that bacteria in phylum Bacteroidetes play a dominant role in HGT events in the human gut microbiome.

**Supplementary Figure 4.**
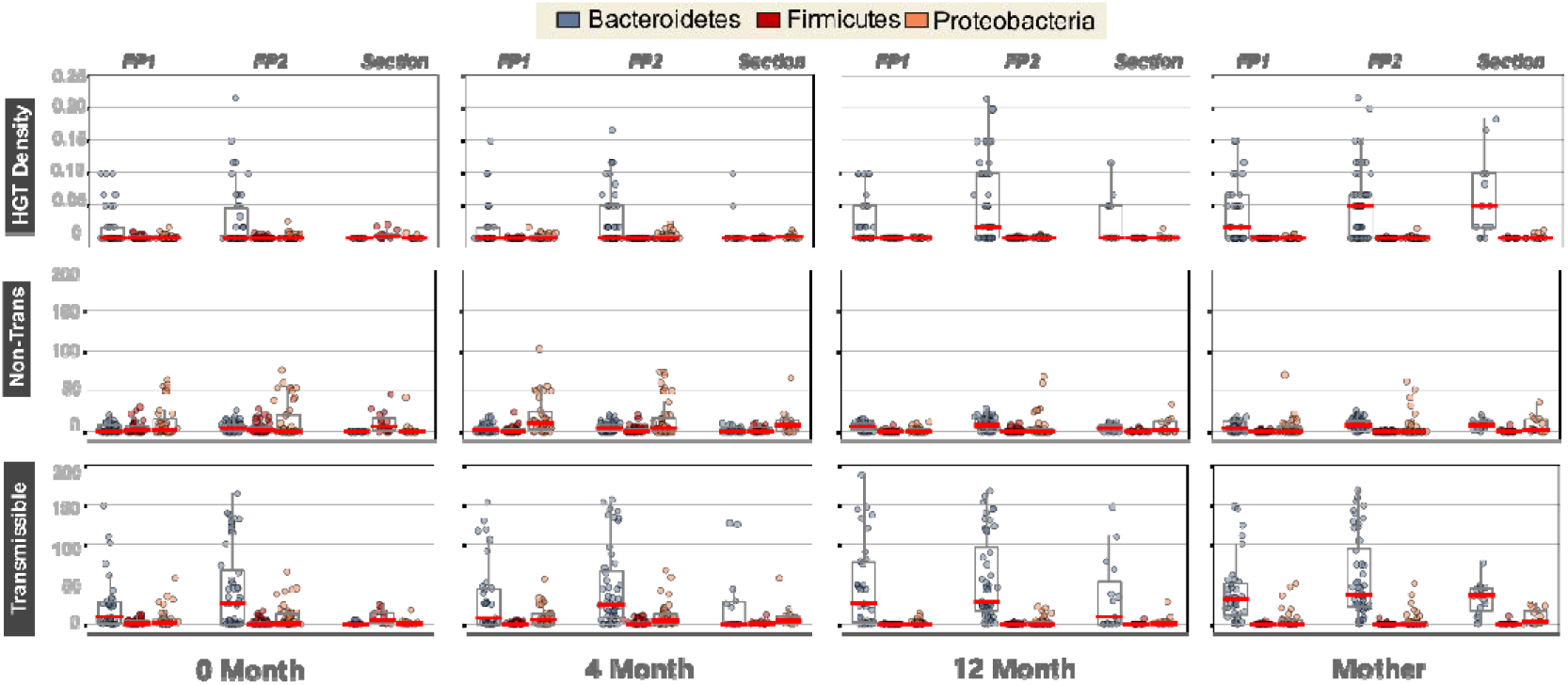
The density of HGT networks in the gut microbiomes of children—distinguished by their age, feeding patterns, and delivery mode—and their mothers. From top to bottom: considering all plasmids, transmissible plasmids only, and non-transmissible plasmids only.

**Supplementary Table 1:** Results of Wilcoxon signed-rank tests comparing HGT events and richness of different types of plasmids among groups of samples.

| Table s1: p values in Wilcoxon signed-rank significance test of HGT event were shown |  |  |  |  |  |  |  |  |  |  |  |
| --- | --- | --- | --- | --- | --- | --- | --- | --- | --- | --- | --- |
|  | fp1 |  |  |  | fp2 |  |  |  | sec |  |  |
|  | b>f | b>p | p>f |  | b>f | b>p | p>f |  | b>f | b>p | p>f |
| new | 0.00163932 | 0.143211222 | 0.15036513 |  | 1.08E-05 | 0.012342746 | 0.024660405 |  | 0.99889114 | 0.87974517 | 0.975069898 |
| 4month | 4.76E-05 | 0.293804272 | 5.08E-06 |  | 1.17E-07 | 0.01025921 | 3.42E-06 |  | 0.07880969 | 0.51564981 | 0.000854492 |
| 12month | 1.06E-06 | 4.31E-06 | 0.000522055 |  | 2.45E-09 | 2.85E-07 | 0.000548128 |  | 0.00464911 | 0.02868652 | 0.004002155 |
| mother | 5.86E-07 | 0.00017742 | 0.000156871 |  | 2.58E-09 | 1.94E-08 | 0.000697123 |  | 0.00110886 | 0.01330566 | 0.008157764 |
| Table s2: p values in Wilcoxon signed-rank significance test of HGT density were shown |  |  |  |  |  |  |  |  |  |  |  |
|  | fp1 |  |  |  | fp2 |  |  |  | sec |  |  |
|  | b>f | b>p | p>f |  | b>f | b>p | p>f |  | b>f | b>p | p>f |
| new | 0.00867726 | 0.005226406 | 0.47164444 |  | 1.45E-04 | 0.000238975 | 0.106669809 |  | 0.99439513 | 0.91014375 | 0.967317877 |
| 4month | 3.69E-03 | 0.068010619 | 2.29E-03 |  | 2.68E-05 | 0.00024249 | 2.10E-04 |  | 0.1425247 | 0.56723284 | 0.007979402 |
| 12month | 2.81E-04 | 2.95E-04 | 0.078649604 |  | 2.01E-06 | 1.49E-06 | 0.063412829 |  | 0.01967975 | 0.02289979 | 0.089856247 |
| mother | 9.02E-05 | 6.82063E-05 | 0.020613417 |  | 2.65E-07 | 2.77E-07 | 0.044040756 |  | 0.00244612 | 0.00288714 | 0.056923149 |
| Table s3: p values in Wilcoxon signed-rank significance test of richness of transmissible plasmid were shown |  |  |  |  |  |  |  |  |  |  |  |
|  | fp1 |  |  |  | fp2 |  |  |  | sec |  |  |
|  | b to f | b to p | f to p |  | b to f | b to p | f to p |  | b to f | b to p | f to p |
| new | 0.00024618 | 0.009568382 | 0.219935618 |  | 2.56E-06 | 0.000494108 | 0.017175956 |  | 0.0041201 | 0.68805957 | 0.012937062 |
| 4month | 5.24E-05 | 0.035374558 | 1.45E-05 |  | 1.65E-07 | 0.000155672 | 0.000142784 |  | 0.15713903 | 0.60984506 | 0.005646655 |
| 12month | 1.72E-06 | 4.60E-06 | 0.000564059 |  | 2.81E-09 | 1.54E-08 | 0.000192333 |  | 0.00506203 | 0.01746176 | 0.067007775 |
| mother | 5.37E-07 | 4.07E-05 | 0.000401094 |  | 2.39E-09 | 1.50E-08 | 7.15E-04 |  | 0.00097871 | 0.00115967 | 9.33E-03 |
| Table s4: p values in Wilcoxon signed-rank significance test of richness of non-transmissible plasmid were shown |  |  |  |  |  |  |  |  |  |  |  |
|  | fp1 |  |  |  | fp2 |  |  |  | sec |  |  |
|  | b to f | b to p | f to p |  | b to f | b to p | f to p |  | b to f | b to p | f to p |
| new | 0.57094328 | 0.032177105 | 0.229628261 |  | 0.221834441 | 0.341343574 | 0.126745576 |  | 0.00331445 | 0.10880943 | 0.040660723 |
| 4month | 0.01842019 | 0.002524963 | 3.51E-05 |  | 0.000910799 | 0.089145168 | 0.000256528 |  | 0.81123541 | 0.00852142 | 0.020169019 |
| 12month | 5.00E-06 | 4.07E-05 | 0.065486818 |  | 1.97E-07 | 5.16E-03 | 0.032566077 |  | 0.01172588 | 0.69886062 | 0.020432514 |
| mother | 3.53E-05 | 2.38E-01 | 0.001440807 |  | 4.14E-09 | 4.82E-04 | 8.51E-03 |  | 9.02E-04 | 0.7527248 | 1.63E-02 |
| f1: exclusively breastfeeding in first four month; no breastfeeding in 12 month |  |  |  |  | b : Bacteroidetes |  |  |  |  |  |  |
| f2: Mix feeding in first four month; any breastfeeding in 12 month |  |  |  |  | f : Firmicutes |  |  |  |  |  |  |
| Sec: c-section born |  |  |  |  | p : Proteobacteria |  |  |  |  |  |  |

### Plasmids expand bacterial gene repertoires

**Supplementary Table 2:**
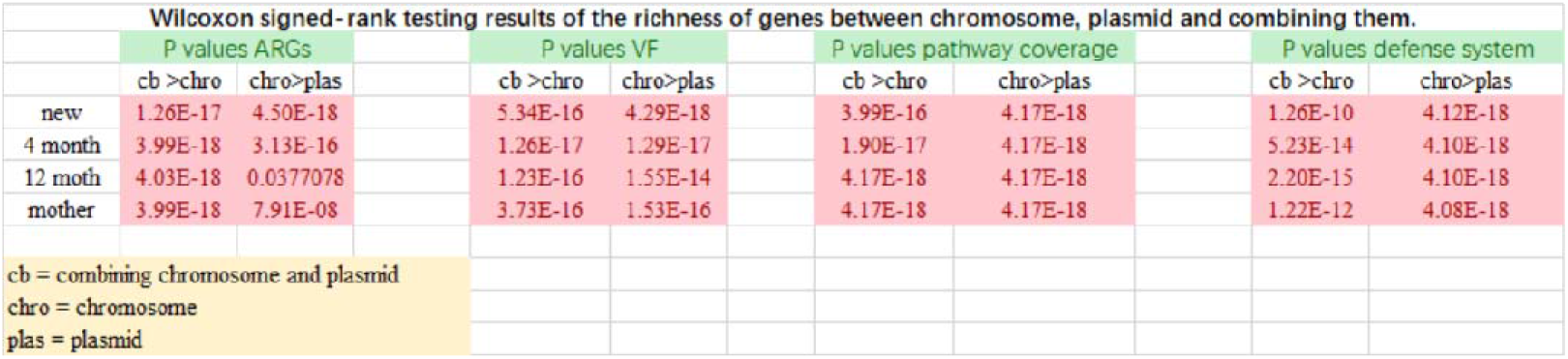
Results of Wilcoxon signed-rank test of gene richness between chromosomes, plasmids, and the combination of the two.

We also assessed the functional capacity of plasmids by annotating their gene repertoires and comparing them to those found on chromosomes. The total amount of chromosomal genes in the bacterial metagenome of our sample was nearly 60 times larger than that of plasmids. Although most plasmid-derived open reading frames (ORFs) were also found on chromosomes, a notable minority—20%—was excusively found on plasmids and may code for plasmid-specific functions (Figure 5a). Overall, this pattern is consistent with the hypothesis that plasmids expand bacterial gene repertoires. The fact that only 32.67% of total plasmid ORFs were successfully assigned functional annotation, compared to ca. 50% annotation rate for chromosome ORFs (Figure 5b). This indicates that plasmids tend to carry more unknown genes than chromosomes. In fact, 25.6% (14,879 of 58,136) of plasmid genes of unknown function were only found on plasmid and never on chromosomal contigs. These are likely to be genes with novel and uncharacterized functions. We also observed that antibiotic resistance genes and virulence factors were found at disproportionately higher rates on plasmids compared to chromosomes (Figure 5b), suggesting that these functional genes are more commonly encoded on plasmids.

**Figure 5.**
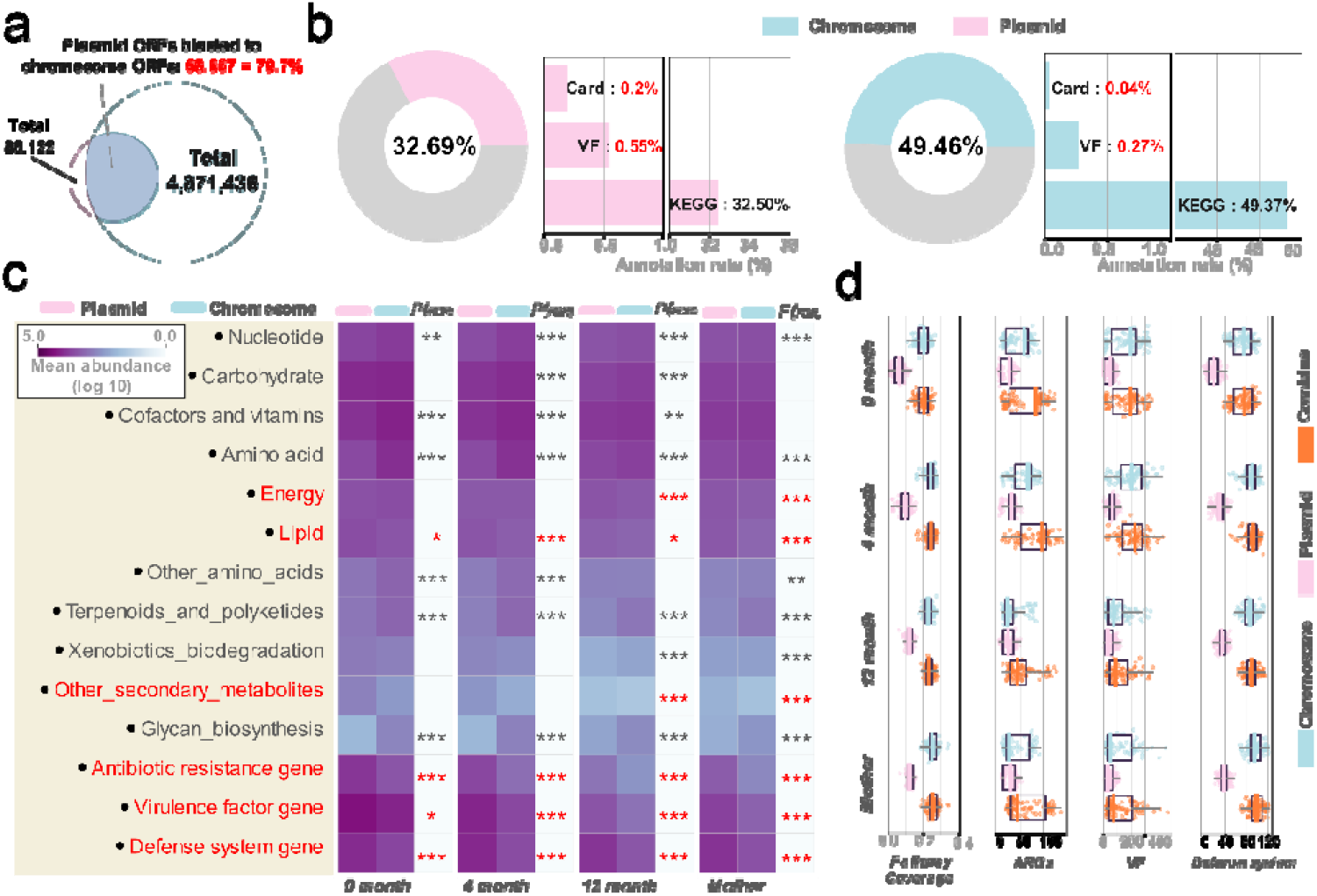
**a**: Open reading frames (ORFs) located on plasmids (pink) were compared to those located on chromosomes (blue) (nucleotide sequence BLAST: coverage >0.8; identity >0.85). **b**: Overall annotation rate (ring diagrams) and database-specific annotation rate (bar diagram) for plasmid and chromosome ORFs. **c**: Total gene abundance of different functional groups. Groups in red were disproportionately present on plasmids compared to chromosomes; groups in gray demonstrated the opposite pattern. Stars indicate statistical significance (Wilcoxon rank-sum test, FDR-corrected, *P ≤ 0.05, **P ≤ 0.01, ***P ≤ 0.001). **d**: The pathway coverage and richness of different functional groups on plasmids (pink), chromosomes (blue), and the non-redundant ORFs combination of the two (orange).

To confirm plasmids’ role in expanding the bacterial gene pool, we analyzed redundancy in gene function between plasmids and chromosomes within each individual in our sample. We found that the gene richness for each functional group was significantly lower in plasmids than in chromosomes, but the coverage of all metabolic pathways was significantly increased when plasmids and chromosomes were combined compared to when chromosomes or plasmids were considered individually (Wilcoxon signed-rank test; Figure 5d, Supplementary Table 2). Similarly, the diversity of antibiotic resistance genes, virulence factors, and defense system genes was significantly higher in the combined genome compared to the chromosome- or plasmid-only genomes (Wilcoxon signed-rank test; Figure 5d, Supplementary Table 2). This suggests that the bacterial gene repertoires were significantly improved by the contributions of plasmids.

Finally, we compared the abundance and richness of genes in different functional groups between plasmids and chromosomes. Plasmids carried numerous metabolism related genes but most of these were also highly abundant on chromosomes (Figure 5c). Notable exceptions, however, were found for genes related to energy metabolism, lipid metabolism, and the biosynthesis of other secondary metabolites. Particularly, lipid metabolism genes were highly prevalent on plasmids in all ages of children and their mothers (Figure 5c). Similarly, genes encoding antibiotic resistance, virulence factors, and components of defense systems were all more abundant on plasmids compared to chromosomes (Figure 5c). Taken together, these data suggest that plasmids play a critical role in promoting bacterial survival and adaptation by providing crucial and potentially novel functions to their hosts.

## Discussion

Here, we investigated the early acquisition and development of the plasmid community in the infant gut—with particular emphasis on the role of plasmids in the larger bacterial community—using a gut metagenomic dataset obtained from a longitudinal cohort study of 98 Swedish children and their mothers. For this, we developed a new bioinformatics pipeline for the isolation and annotation of plasmids from metagenomic data. With these data, we constructed a phylogenetic tree of plasmids based on their conserved backbone replicon genes, and developed a novel strategy for the identification of potential bacterial hosts based on genomic similarity to known plasmids. Our findings highlight the importance of plasmid biology for the gut microbiota in early life.

One important finding was that plasmid assemblages may be more sensitive than their bacterial host communities to environmental variation (Figure 1c). Although unexpected, this is consistent with previous studies reporting that, due to their higher diversity and copy number^2^, plasmid-encoded genes evolve faster than chromosomal genes^20,21^. Plasmids may therefore be able to adapt faster to changing environmental conditions. This could mean that, compared to bacteria, plasmid colonization of the infant gut is less influenced by inheritance from the mother, since the newborn gut environment differs from that of the mother in significant ways (e.g., bacterial density, oxygen concentration, host modulation) (Figures 1d and e). In this way, plasmids might serve as mechanisms for rapid adaptation to environmental variation. It is possible that, upon arrival in the newborn gut, bacterial hosts inherited from the mother^57,58,59^ rapidly exchange and selects new plasmids that help the community adapt to the new gut environment. Nonetheless, it cannot be excluded that the better resolution seen on the plasmid community composition is not a reflection of individual strain-level specificity between which cannot be accounted for with precision using short reads metagenomics.

One of the key findings of this study is that plasmids are enriched in the infant gut in the first four months of life (Figure 2b). This makes sense in light of previous studies demonstrating that plasmids play an important role in bacterial evolution and adaptability by transferring beneficial traits within and between species of bacteria, positively contributing to host fitness^8^. The establishment of the early gut microbiome is dominated by stochastic processes that favor rapidly proliferating bacteria^60^. The strength of the competitive pressure in such an environment should favor rapid adaptation, likely increasing the importance of mobile genetic elements carrying adaptive traits. It is plausible that some bacteria randomly seeded at birth may lack specific colonization factors and may therefore depend on transmissible vectors for these important functions, thus explaining the increased abundance and diversity of plasmids observed at 4 months of age. Then, as the bacterial community matures, it moves toward a more functionally uniform^60^ and constrained microbiota, with essential traits shared at the phylum level^61^, and moves away from an extensive pan-genome of mobile elements. This would explain the reduced diversity of the plasmid community that we observed at 12 months of age, as the bacterial assemblages begin to resemble those of adults (Figure 2b). One possible factor in this decline could be the fitness cost of plasmids, which is the main limit to their persistence in bacterial populations^19,23,62^. As children grow and the functionality of the gut ecosystem gradually stabilizes, the community composition become more fixed and a network of mutualisms is developed^57,36,60^, meaning that the benefits of plasmids for bacteria—i.e., rapid adaptation to stochastic changes—may start to be outweighed by their fitness costs, promoting plasmid loss. Another indisputable factor is breastfeeding, as an earlier study showed that the maturation of the infant gut microbiome was directly associated with the cessation of breastfeeding^36^. Indeed, we observed that, with respect to plasmid diversity, 12-month-old children who were breastfed in any amount did not differ from newborns and 4-month-olds, but 12-month-olds who were not breastfed demonstrated a significant reduction. This finding supports previous research on the importance of maternal breast milk for the infant gut microbiome and mobile genetic elements^63,64,65^, but, in general, this effect is very poorly understood. Specifically, it is unknown if breast milk is the main source or driver of the infant gut microbiome, and how this might compare to other types of maternal-infant transfer.

Mobility genes are critically important for plasmids because they facilitate horizontal transfer among bacterial cells^5^. Here, we also found that plasmid mobility can impact prevalence (Figure 2a), corroborating previous suggestions that a sufficiently fast transfer rate can compensate for fitness costs and plasmid loss^12,66^. Moreover, we found that mobility genes are not distributed randomly in the plasmid population: we identified a clear phylogenetic signal (relatedness) in the replicon groups in which these genes were found (Figure 2a).

In the literature, plasmids are well known from the intensively studied bacteria in family Enterobacteriaceae, especially *Enterococcus, Escherichia*, and *Klebsiella*^67,68,69,70^, which has led to the overrepresentation of these examples in databases such as PLSDB (Figure 3a)^49^. However, we found that the main bearers of plasmids—particularly transmissible plasmids—in the human gut are *Bacteroides* and *Bifidobacterium*. Indeed, by its reliance on a biased database, it is likely that our method still underestimates the plasmid diversity of these two bacteria (Figure 3d). Similarly, a previous study reported that multiple interspecies and intergenus HGT events are facilitated by *Bifidobacterium* and *Bacteroides*^30^. This concordance supports the reliability of our host linking method, which may serve as an important tool for identifying plasmid hosts in metagenomes of the gut environment.

Consistent with a report that HGT is biased towards more closely related individuals and species and occurs more rarely between distant relatives^71^, we found that putative plasmid-mediated HGT decreased with the phylogenetic distance between bacteria. In children of all ages and their mothers, Bacteroidetes had the highest frequency of potential plasmid HGT (Figure 4a, Supplementary Figure 2a), because this genus contained the highest abundance of transmissible plasmids (Figures 3a, 3d, 4a and Supplementary Figure 2a). This finding supports previous results that almost half of the HGT events in the human gut are mediated by members of phylum Bacteroidetes^3072^. Likewise, our finding that Proteobacteria and Firmicutes have the most widespread HGT networks (Figure 4a, Supplementary Figure 2a) makes sense given previous research revealing that plasmids are very prevalent in members of these phyla^8^.

This work highlights the role of plasmids in expanding the gene repertoires of their bacterial hosts. We detected a larger number of new and unknown genes on plasmids than on chromosomes, which could possibly be due to the typically high copy number of plasmids, which promotes a higher mutation rate^58^. Moreover, of the genes that were found on both plasmids and chromosomes, some were found in higher abundance on plasmids, which supports previous studies reporting that plasmid genes usually have higher gene dosage effects than chromosomal genes^19^. It could also be that certain genes have a stronger fitness effect for the host bacterium when present on a plasmid compared to on a chromosome.

The methodological approach in this work mainly relies on our previously developed plasmid analysis pipeline, with some improvements in performance and reliability. Still, though, only a limited number of plasmids were identified, which we believe is likely to be a function of sequencing bias. In the future, efforts should be made to improve plasmid sequencing technology in order to isolate more plasmid genomes and enable more comprehensive analyses of plasmids in the gut. Notably, although our method was successfully able to identify the hosts of many plasmids based on their genetic similarity to examples in the PLSDB database, we were inherently constrained by the limitations of the database: only 12.8% of the plasmids in our sample were successfully linked to potential bacterial hosts. Furthermore, because it relies on previously published knowledge, this method cannot be used to detect novel plasmid-carrying bacteria. Future work to improve detection algorithms and expand the plasmid database will help in this respect. Although our current understanding of gut plasmid biology is still limited, this study provides strong evidence that plasmids play an indispensable role in the colonization and development of the gut microbiota in early life.

## Conclusion

In this work, we constructed a catalog of plasmids present in the human gut with the goal of providing, for the first time, baseline information about the acquisition and development of the gut plasmidome in early life. Plasmids in the infant gut were extremely diverse and, in some respects, better reflected environmental changes than species level community composition. Plasmid abundance and diversity was particularly high in the first four months of life and decreased with maturation of the gut microbiota. Importantly, we discovered that *Bacteroides* and *Bifidobacterium* serve as major hosts of transmissible plasmids in the early human gut microbiota; this is especially relevant because much of the existing literature on plasmids focuses on the host bacteria *Escherichia* and *Klebsiella*. Finally, we found that gut plasmids play a substantial role in the expansion and abundance of the gene repertoires of their bacterial hosts.

## Author Contributions

JN and SJS designed the study; WLH processed the bioinformatic and statistical analysis, produced the figures, and wrote the manuscript; JR gave recommendations for bioinformatic and statistical analyses; JN and SJS reviewed and provided suggestions for bioinformatic analyses. All authors commented on and approved the final manuscript.

## Acknowledgments

This research was partly funded by the European Union’s Horizon 2020 Research and Innovation Program under grant agreement No. 818431 (SIMBA, Sustainable Innovation of Microbiome Applications in the Food System). WH is funded by the China Scholarship Council (CSC) No. 201806370229. We are grateful for the work of the Swedish cohort project (ERP005989).

